# Tracking cancer dynamics from normal tissue to malignancy using perfect N- and T-gene expression markers

**DOI:** 10.1101/2024.11.04.621130

**Authors:** Gabriel Gil, Rolando Perez, Augusto Gonzalez

**Affiliations:** Institute of Cybernetics, Mathematics and Physics, Havana; Center for Molecular Immunology, Havana

**Keywords:** N- and T-genes, Carcinogenesis, Somatic evolution, Clonal evolution

## Abstract

Common knowledge states that the spontaneous somatic evolution of a normal tissue may lead to a tumor. Once the tumor is formed, it naturally evolves towards a state of higher malignancy. On the other hand, perfect gene expression markers for normal tissue and tumor—the so-called N-genes and T-genes—were recently introduced. We join these two pieces of knowledge in order to argue that: 1) Only N-markers participate in the spontaneous dynamics of a normal tissue. The number of active markers decreases as the tissue approaches the transition point where it becomes a tumor. 2) Only T-markers participate in the spontaneous dynamics of tumors. The number of markers increases as the tumor becomes more malignant. 3) Both sets of genes are connected by the so-called NT-genes, i.e., genes that are simultaneously N- and T-markers. They should play a crucial role at the transition point and, possibly, when the tumor is exposed to a drug or therapy. 4) The pathways or mechanisms protecting the normal tissue from becoming a tumor may be described by a small perfect panel of N-genes. 5) The pathways or mechanisms guiding the evolution of tumors in a tissue may be described by a small perfect panel of T-genes. We illustrate the above statements with the analysis of expression data for prostate adenocarcinoma, one of the most heterogeneous tumors. In this case, there are about 1000 N-genes and 6000 T-genes, and the perfect N- and T-panels contain 11 and 8 genes, respectively. Additionally, we provide examples from lung adenocarcinoma and liver hepatocarcinoma.

## 1. Introduction

The phenomenological multi-step model of cancer states that accumulated genetic and epigenetic changes in a normal tissue may lead to a tumor [1]. Attempts have been made to formulate a kind of multi-step model at the gene level. One example is the Vogelstein model of colon cancer [2]. However, no detailed model exists for an arbitrary cancer localization. We understand that oncogenes should be activated and tumor suppressor genes deactivated, but nothing more. Which are these genes? How many should be activated or deactivated in order to trigger the transition? These questions do not have a definite answer, in general.

On the other hand, once the tumor is formed, it evolves according to the laws of clonal evolution [3]. In the differential expression approach to gene expression analysis, both processes—carcinogenesis and clonal evolution of tumors—are treated as a single event. The relevant genes, ranked only by differential expression values, are not linked to one process or the other.

Consider, for example, the *EPHA10* gene in prostate adenocarcinoma (PRAD). In Fig. 1a we show expression histograms, constructed from TCGA data [4,5], in normal and tumor samples. The gene is overexpressed in tumors. It may be classified as an oncogene. But where does it play a role: in the process of tumor formation or in its evolution once formed?

**Figure 1.**
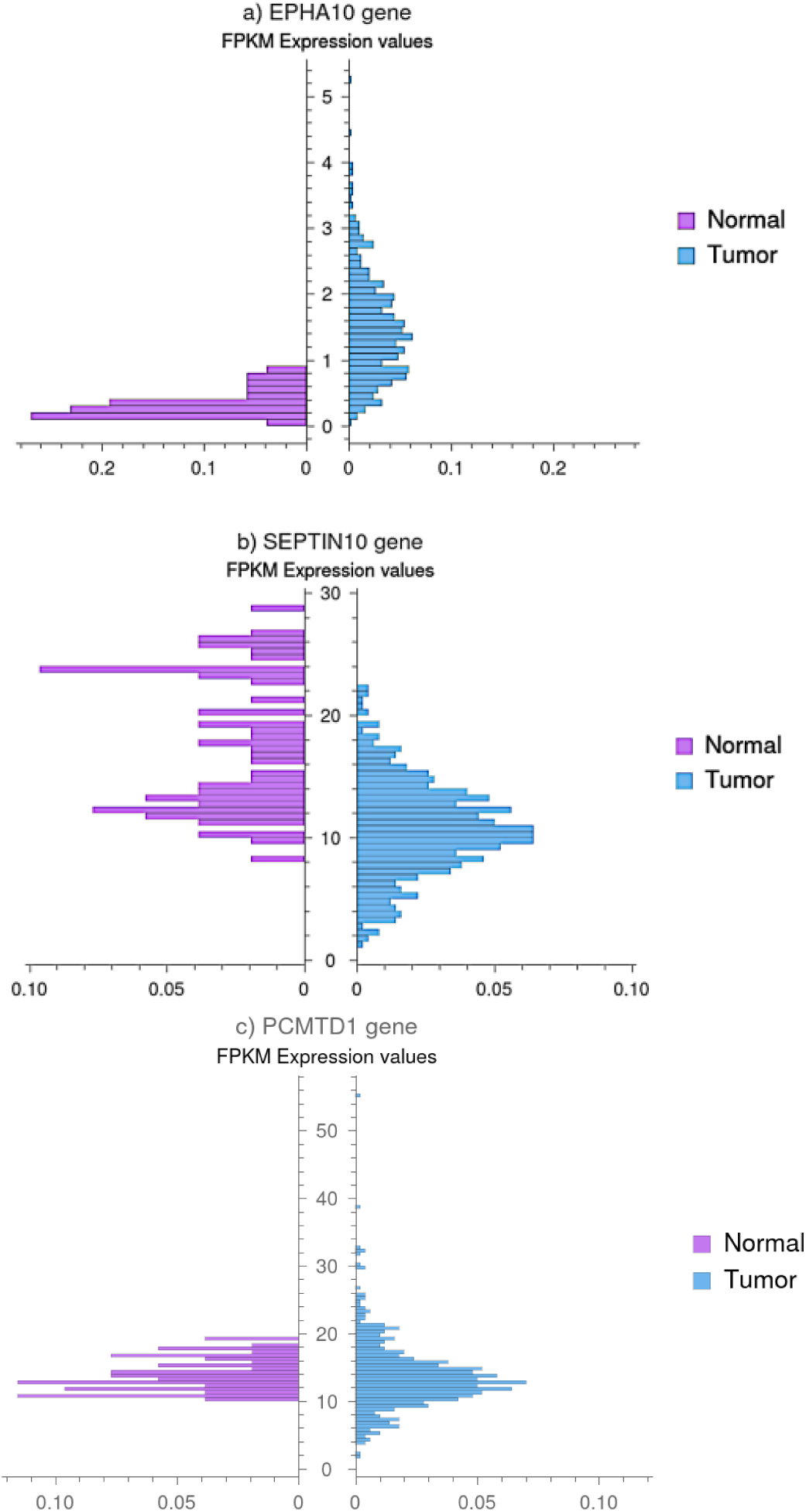
Paired probability histograms of FPKM expression values in PRAD. a) The EPHA10 gene, a tumor-above gene. b) The SEPTIN10 gene, which can be included either in the normal-above group or in the tumor-below group. c) The PCMTD1 gene, which belongs to the tumor-outside category, exhibiting expression intervals both above and below the normal range.

In the present paper, we formulate a new paradigm of gene expression analysis based on genes that are perfect tissue markers [6]. These genes show expression intervals characteristic only of normal or tumor samples, respectively. We call them N-genes and T-genes.

The *EPHA10* gene shown above is an example of a tumor-above gene. We may explicitly state that it plays a role in the clonal evolution of tumors. More precisely, the expression of this gene does not experience significant variation during carcinogenesis. Once the tumor is formed, however, it may reach high expression values, becoming in this way a tumor marker.

We use a kind of ergodic principle [7] to infer time evolution from static gene expression data. In this way, we show that N- and T-genes are also useful in describing dynamical processes such as carcinogenesis [8] or the clonal evolution of tumors. Perfect gene panels, constructed from N-genes and T-genes [6], may be used to label the way somatic evolution in carcinogenesis or clonal evolution of tumors proceeds. The emerging picture for carcinogenesis is similar to the multi-step model of cancer [1].

The plan of the paper is as follows. In the Methods section, we introduce a few sentences on the data and code used. For the sake of consistency, we also recall the definitions of N-, T-, NT-, and O-genes. In the Results and discussion section, we show how the number of activated N-genes is reduced in the process of somatic evolution of a normal sample. Clonal evolution of tumors, on the other hand, is shown to be related to a continuous increase in the number of T-markers. Next, a conceptualization of the transition from normal tissue to tumor is presented. The introduction of perfect gene panels, according to Ref. [6], allows us to label the evolution of normal and tumor tissues. Finally, at the end of the section we discuss common driver genes in light of the present framework. The paper ends with a Concluding remarks section.

## 2. Methods

### 2a. Data and code

Gene expression data for normal tissue samples and tumors were obtained from The Cancer Genome Atlas public network (TCGA, https://portal.gdc.cancer.gov/) [4]. We illustrate the analysis with data for prostate adenocarcinoma (PRAD) [5], lung adenocarcinoma (LUAD) [9], and liver hepatocarcinoma (LIHC) [10].

These are expression data for tissue samples. In principle, the analysis could be performed on the basis of single-cell RNA-seq data [11]; however, the diversity of cells makes this more complicated. Thus, we use traditional sample data. N-genes and T-genes are understood as tissue markers.

The code is very simple: finding the maximum and minimum of a set of values, building histograms, comparing two sets to find tumor-only expression intervals, comparing two lists to find common genes, etc. It may be obtained from the authors upon request.

### 2b. N-genes, T-genes, NT-genes, and O-genes

There is a simple abstract picture in which the concepts of N- and T-genes become apparent. Indeed, the N- and T-attractors are far apart in gene-expression space. Using the 60,000 reported genes in TCGA data and log-fold coordinates to compute the Euclidean distance [12,13], this distance appears to be around 92.7 in PRAD [12,14], for example. Let us introduce a coordinate *x* along the axis joining the centers of the N- and T-attractors. Values of *x* around 0 are characteristic of normal samples, whereas values around 92.7 are characteristic of tumors.

The *x* coordinate is a metagene [15], which contains the contribution of many genes. Thus, one may qualitatively understand that there should be many coordinates (genes) contributing to the *x* variable, with expression intervals characteristic only of normal samples or of tumors. We call them N-genes and T-genes, respectively. In the same way, coordinates “orthogonal” to the *x* axis may have expression intervals accessible to both normal samples and tumors. We call them O-genes (O for others).

This simple and abstract argument for the existence of N- and T-genes may be easily verified in the expression data. Consider, for example, the aforementioned *EPHA10* gene in PRAD (Figure 1a). It is an example of a T-gene. The FPKM normal and tumor intervals for expression are (0.04, 0.88) and (0.08, 5.25), respectively. Thus, there is an interval, (0.88, 5.25), characteristic only of tumor samples. We discretize the expression and write *e* = 0 for normal and tumor samples in the common interval, *e* = 1 for tumor samples with expression above 0.88, and include the gene in the “tumor-above” group. Notice that *e* = 1 for this gene is an indication of a tumor, whereas *e* = 0 for all normal and for some tumor samples. When *e* = 1, we say the gene is T-activated.

On the other hand, for N-genes there is a distinct interval for normal samples. In this case, we write *e* = 0 for expression values of both tumor and normal samples in the common interval, and *e* = −1 for normal samples in the normal-only characteristic interval. In Figure 1b we show paired histograms for the *SEPTIN10* gene in PRAD. As is apparent, we may include this gene in the normal-above group. Values of expression above 22.26 seem to be incompatible with the tumor state and are characteristic of normal samples.

Notice, however, that this latter gene also exhibits a tumor-only interval, in which the expression is below 8. Thus, it behaves as a normal-above gene when the expression is above 22.26, or as a tumor-below gene when the expression is below 8. The class of genes with this property is called NT-genes because they belong to both the N- and T-genes subgroups. As will become apparent below, if *SEPTIN10* is a normal marker of a given sample, it should be deactivated during carcinogenesis. Its expression should change from −1 to 0. Once the tumor is formed, it may be activated as a T-marker, with its expression changing from 0 to 1.

We introduce the “normal-above,” “normal-below,” and “normal-outside” categories, along with similar categories in the T-genes group. The NT-genes come in two variants: tumor-above–normal-below and tumor-below–normal-above.

The non-differential character of our classification is best seen in the “outside” sets. For example, the tumor-outside group is composed of genes that are simultaneously tumor-above and tumor-below, i.e., there are expression intervals for tumor samples above and below the normal region. The common region for tumor samples and some normal samples is sandwiched between the external intervals. An example is the *PCMTD1* gene, whose histogram is shown in Fig. 1c. This gene would never arise in a differential expression approach, although it has been identified as a fusion gene in PRAD [16]. Deregulation of this gene in expression data could be related to fusion events.

The N- and T-genes are markers of the states. But N- and T-genes may also be understood as holders or anchors of the states, in the sense that a set of activated N-genes, for example, indicates that we are in the vicinity of the normal attractor. To move the sample away from the attractor, we must first inactivate the anchors. We will return to this point below when analyzing the evolutionary dynamics.

In principle, for sufficiently large sets of samples, the numbers of N- and T-genes in a tissue should reach saturation values. But for finite sample sets, these numbers are set-dependent. Indeed, to include a gene in a group, we require the normal-only or tumor-only intervals to be populated by enough samples, and we establish a statistical significance criterion by means of a simple exact Fisher test [17]. Let *n* be the number of samples in the normal-only or tumor-only gene expression interval. Requiring that *n* not be the result of chance, with a p-value of 0.01, the test leads to:

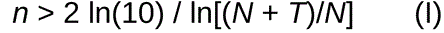

or

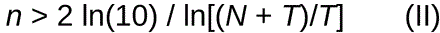

for N-genes and T-genes, respectively. In Eqs. (I) and (II), *N* and *T* correspond to the numbers of normal and tumor samples. In the TCGA data for PRAD, *N* ∼ 50, *T* ∼ 500; thus, the tests lead to *n*/*N* > 4% and *n*/*T* > 9%.

In this paper, we use the stricter thresholds of 5% and 10% to define N- and T-genes in PRAD. We found 1097 and 6138 genes, respectively. Three hundred sixteen genes are in the NT category. Notice that the number of perfect marker genes is around 7200. The rest of the 60,000 genes in the TCGA data are either not expressed in the tissue or are O-genes. This represents a considerable reduction of the expression space to be analyzed. Notice also that there are many T-genes and far fewer N-genes. This is a consequence of the higher disorder of tumors [18,19].

Let us summarize. There are T-genes, whose discretized expression may take the value *e* = 0 (the common interval for N and T samples) or *e* = 1 (an interval characteristic only of tumors). In this latter case, we say the gene is T-activated and the sample is, of course, a tumor. In the same way, there are N-genes, whose expression may take values *e* = 0 or *e* = −1; in the latter case, we say the gene is N-activated. This can occur only in a normal sample. Besides, there are NT-genes, whose expression may take values *e* = 1, 0, and −1. That is, they may transit from N-activation to T-activation and vice versa. Additionally, there are genes whose expression is always *e* = 0. Their expression intervals are accessible to both normal samples and tumors. These are the O-genes.

## 3. Results and discussion

### 3a. N-markers and the somatic evolution of a normal sample

We first argue that in the spontaneous somatic evolution of a normal sample, only the N-markers are important. Indeed, let us imagine that we use a discrete expression model to describe the dynamics, as in Ref. [20]. In this model, the O-genes always take the value *e* = 0 and do not participate in the dynamics. With regard to the T-genes, they cannot be permanently activated to *e* = 1 within a normal state. Thus, the relevant dynamics is the dynamics of N-markers.

In the differential expression approach to expression analysis, normal samples are used only to set reference values for gene expression. However, normal samples are ideal for the study of carcinogenesis. They are representative of normal tissues in different stages of somatic evolution.

We show in Fig. 2 a reduced-dimension representation of the PRAD data. Principal component analysis (PCA), along the lines of Ref. 13, is shown in Figure 2a. The center of the normal cloud of samples is at *x* = 0, whereas the center of the tumor cloud is at *x* = 92.7. In Figure 2b we use the same *x* axis, but the ordinate is the number of activated N-markers in the 52 adjacent normal samples of the data. Notice that this number ranges from 1 to about 600. Samples closer to the center of the normal attractor exhibit healthy conditions associated with larger numbers of activated N-genes (500 or more), whereas the number is reduced in the course of somatic evolution as the samples approach the tumor attractor. It is remarkable that a single activated N-gene may maintain the normal-state condition, even after more than 500 genes have been deactivated.

**Figure 2.**
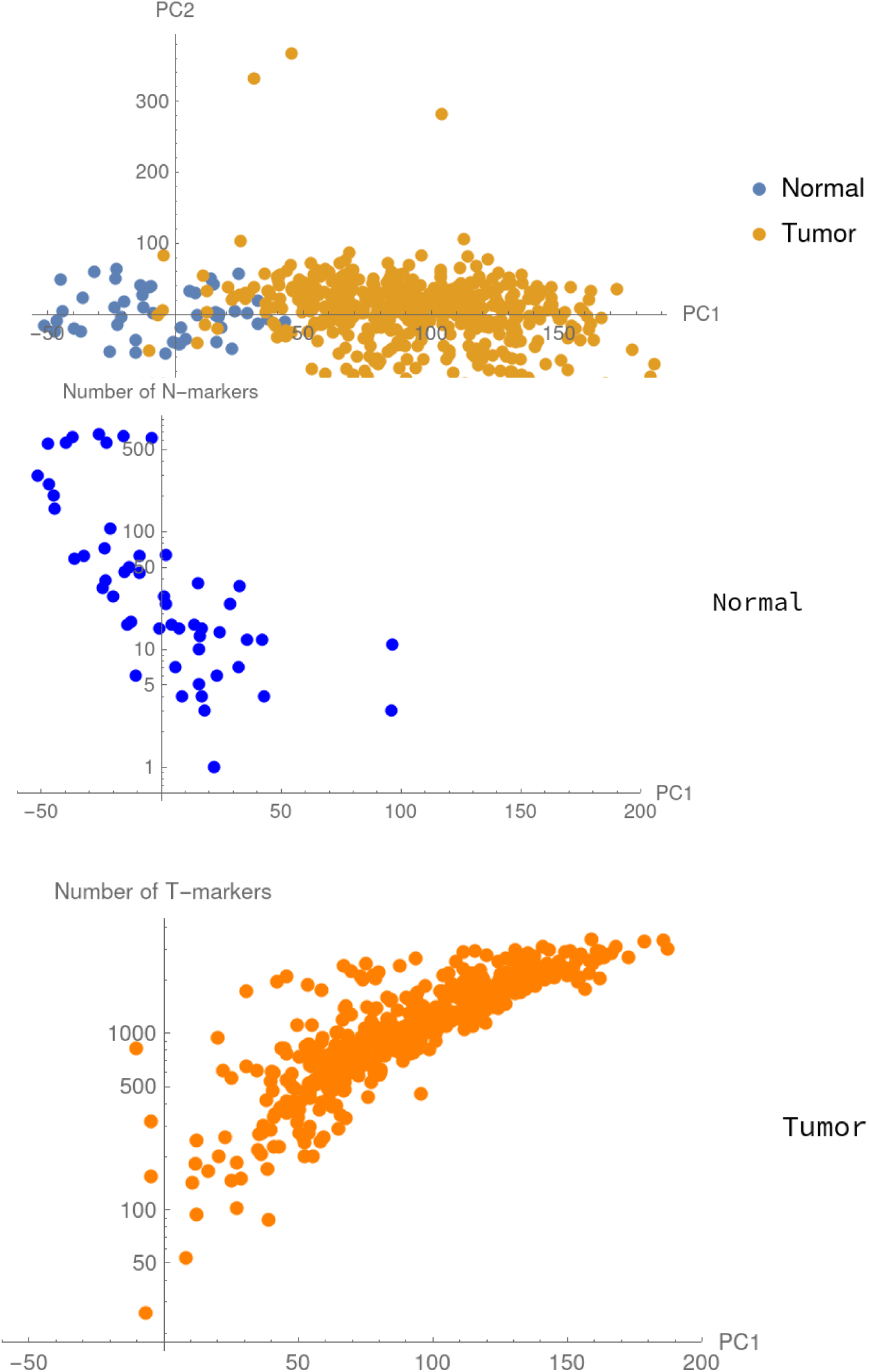
Two-dimensional representation of PRAD data and the number of N- and T-markers. a) Principal component analysis showing clouds of normal and tumor samples. b) Number of N-markers as a function of the PC1 coordinate. c) Number of T-markers as a function of the PC1 coordinate.

We know that distances are not properly represented in PCA diagrams. We correct this distortion in Figure 3a, where we use as the *x*-axis the distance in gene-expression space from the normal sample to the center of the tumor attractor. The functional dependence becomes more apparent. We draw a solid line only to guide the eyes, but it is apparent that the tendency is toward zero activated N-genes as the transition to the tumor is approached.

**Figure 3.**
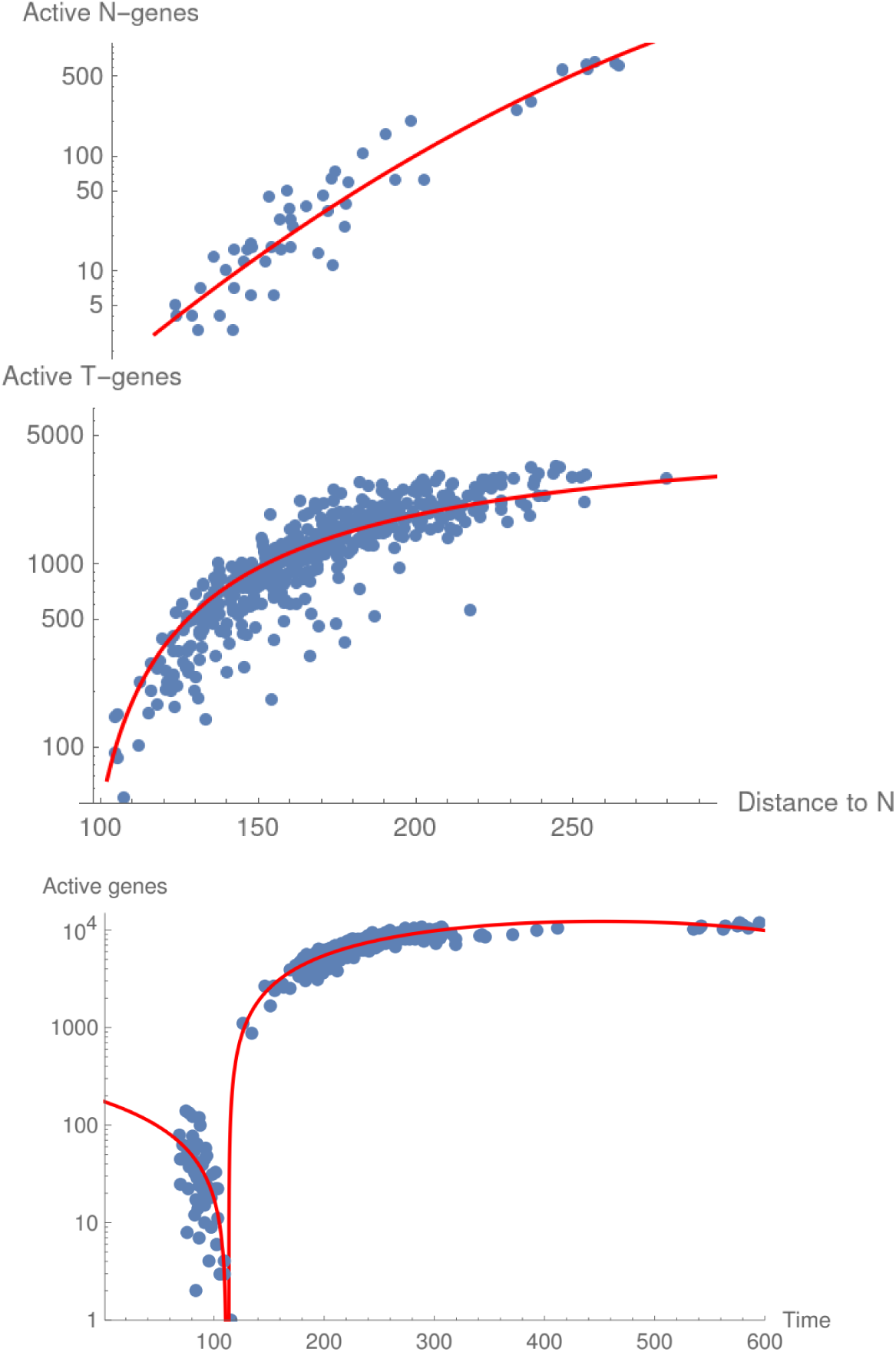
Correction of distances in Figs. 2b and 2c. a) Number of N-markers as a function of the distance from the normal sample to the center of the tumor attractor. b) Number of T-markers as a function of the distance from the tumor sample to the center of the normal attractor. c) Schematic representation of the transition. Figs. 3a and 3b are combined, and the *x*-axis is transformed to time using the ergodic principle. Red dashed lines are drawn as a guide to the eye.

By using a kind of ergodic argument [7], we may interpret Figures 2b and 3a as the schematic somatic evolution of normal samples, which continuously lose N-markers as they approach the border of the basin of attraction of the N-attractor, becoming in this way closer to the T-attractor. We shall use the same argument in the next sections. Let us stress that in the data, corresponding to measurements at a single time point, the information on which N-markers were initially activated in a healthy sample is lost. What we measure are the remaining N-markers, not the deactivated ones.

In Supplementary Figures 1a and 1b, we show results similar to Fig. 3a for lung adenocarcinoma (LUAD [9]) and liver hepatocarcinoma (LIHC [10]). The numbers of N-genes in these tissues, with the same threshold of 0.05, are 492 and 581, respectively.

### 3b. T-markers and the clonal evolution of tumors

In the same way, we may argue that only T-markers are relevant in the spontaneous evolution of tumors, and use them to examine the set of tumor samples and infer the dynamics of clonal evolution. To this end, we plot in Figure 2c the number of activated T-genes as a function of the PC1 coordinate. This number ranges from 25 to around 3300 in PRAD. The figure shows tumor samples exhibiting very different stages of clonal evolution. Those with a relatively small number of activated T-genes are closer to the normal attractor. They probably correspond to recently formed tumors. Evolved tumors show higher numbers of activated T-genes. Similar to the figure for normal samples, we use a corrected distance from the tumor sample to the center of the normal attractor to improve the functional dependence and generate Figure 3b. The tendency is toward zero activated T-genes at the transition.

Supplementary Figures 2a and 2b show similar results for LUAD and LIHC, respectively. The number of T-genes in these tumors, with the same threshold of 0.1, are 17037 and 17351, respectively.

In summary, Figures 2c and 3b show that clonal evolution is characterized by a continuous increase in the number of T-markers as the sample moves away from the N-attractor and approaches the central area of the T-attractor. Unlike somatic evolution of normal samples, where initially activated genes are lost, a gene expression measurement in a tumor sample shows all of the T-markers activated at a given time.

### 3c. Conceptualizing the transition

Recall the diagram in Figure 2a. The normal and tumor attractors are separated by a low-fitness barrier [12]. The transition between attractors is discontinuous, according to concepts from statistical physics [21,12].

We may draw the following schematic picture for the transition. The initial condition is a small portion of the tissue near the center of the normal attractor (a healthy normal sample). Somatic evolution consists of small jumps inside the basin until it reaches a point somewhere near the border. Then a large jump toward the tumor basin of attraction occurs [22]. Subsequently, small jumps within the tumor basin represent clonal evolution.

In terms of N- and T-genes, first the N-genes holding the normal state should be gradually inactivated. When all of the holders are down, the transition occurs and new T-markers arise. A schematic picture is presented in Figure 3c. The *x*-axis in that figure is constructed from the distances in Figures 3a and 3b, transforming distances to time with the help of the aforementioned ergodic principle. The transition is represented as a point, although it could also be an interval or region where N-markers and T-markers coexist. Thus, to describe the dynamics at the transition point, we must include both N- and T-markers and their regulation networks.

The number of activated N-genes in the normal sample is reduced as a result of perturbations. Perturbations to the normal state come in the form of somatic mutations [23], epigenetic alterations [24], or external influences [25]. They may provoke deregulation cascades [20] in the regulation network.

Assume the system is in a normal state and a perturbation occurs. The attractor is a stable state and may support small perturbations. If the cascade reaches a subset of the activated N-genes, it may modify them, moving their expressions to the interval common to N and T samples—that is, reducing the expression to *e* = 0. The reached genes stop acting as holders of the N state, and a step toward the tumor is taken. In the process, part of the information about the initial normal state (the initially activated N-genes) is lost. Notice that the expression of a gene cannot be moved toward T-activation intervals if there remain activated N-genes holding the normal state.

Thus, the first cascades prepare the transition by inactivating most of the N-genes. There should be a cascade that actually realizes the transition, where no N-genes are left and a set of activated T-genes marks the new tumor. Near the transition point, both N- and T-deregulation networks are important, and the NT-genes, which may change their activation state, could play a crucial role connecting both networks.

The idea of cascades in carcinogenesis is also supported by gene expression data. Indeed, the data show blocks or bunches of N-genes with the same expression profile in normal samples. That is, all genes in a block are simultaneously activated in the same normal samples. The largest block in PRAD contains 212 genes. Then, in the somatic evolution of the normal tissue, there could be a single event in which the tissue loses 212 markers. This situation recalls the multistep model of cancer [1]. We present in Table I the frequency and number of genes for some of these blocks.

**Table I.** Properties of panel and block genes in the PRAD data. The main five block genes and the genes comprising the perfect N- and T-panels are included.

| Block N-genes | Frequency in normal samples | Number of genes in the block |
| --- | --- | --- |
| Block 1 | 0.13 | 212 genes |
| Block 2 | 0.11 | 33 genes |
| Block 3 | 0.05 | 28 genes |
| Block 4 | 0.11 | 26 genes |
| Block 5 | 0.15 | 20 genes |
| Perfect panel of N-genes | Type and Frequency in normal samples | Reported protective role |
| SEPTIN10 | N-above, T-below, 0.30 | Ensures accurate cell division (through its interaction with BUB3) and facilitates the death of cells with damaged microtubules [26]. SEPTIN10 acts as a dual caretaker and gatekeeper. The protective function is likely dependent on the specific protein environment of a healthy prostate cell. If that environment changes—for example, if a tumor does form—SEPTIN10's role could be subverted to support cancer progression [27], as seen in other tissues like the liver. |
| CA14 | N-above, 0.19 | Regulates pH balance and, as part of a normal metabolic program, helps maintain a cellular environment that is hostile to cancer initiation [28,29]. Loss of CA14 expression is a key step in prostate cancer development and progression [30]. |
| TES | N-above, T-below, 0.15 | Its protein product helps maintain normal cell structure and adhesion [31]. It can activate stress-response pathways like p38 to eliminate potentially dangerous cells [32]. It is a key "gatekeeper" in the prostate epithelium [33]. |
| PARM1 | N-above, T-below, 0.15 | Appears to function as an androgen-repressed, pro-survival factor. Its primary role in prostate cells may be to help them survive periods of low androgen. This could be seen as a protective mechanism for the cell, but it's one that cancer cells can exploit to their advantage, promoting immortalization and proliferation [34,35,36]. |
| APOBEC3C | N-above, T-below, 0.13 | Functions as a tumor suppressor in prostate cells [37]. Its expression maintains normal growth control, suppressing proliferation, migration, and invasion . Its loss is an early step that removes a barrier to cancer development. It is one member of a family of enzymes. While it protects, other family members (like APOBEC3B) are potent mutators that drive |
|  |  | cancer progression, heterogeneity, and therapy resistance [38] when they are "unleashed" by the loss of regulatory proteins like SYNCRIP [39]. |
| ENSG00000224984 | N-above, 0.13 | Novel transcript. lncRNA |
| TMT1A | N-above, 0.11 | Functions as a multi-faceted tumor suppressor by: 1) Directly inhibiting cancer cell proliferation and migration (potentially via p53) , 2) Shaping a favorable immune microenvironment by suppressing tumor-promoting M2 macrophages and enhancing T cell activity, and 3) Potentially regulating cell-cell communication through its RNA methyltransferase activity [40]. The significant downregulation of TMT1A in prostate adenocarcinoma tissues is entirely consistent with it being a tumor suppressor whose loss is selected for during cancer development. This mirrors its behavior in lung cancer and melanoma [41]. |
| SLC18A2 | N-above, 0.09 | Essential for normal neuroendocrine cell function in the prostate. By regulating the release of signaling molecules, it helps maintain tissue homeostasis and suppresses aberrant epithelial proliferation [42]. It acts as a gatekeeper of the normal prostate microenvironment. It is specifically and comprehensively silenced in prostate cancer through early, concurrent genetic loss (LOH) and epigenetic hypermethylation [43,44]. This silencing removes its protective influence, creating a permissive environment for malignant transformation. |
| SNORD104 | N-below, T-above, 0.07 | There is not current evidence for a protective effect in prostate, although other SnoRNAs have been shown to promote proliferation and migration in prostate cancer [45]. The emerging picture from endometrial cancer suggests it is an oncogene that could potentially drive prostate cancer progression through PARP1 regulation [46]. |
| RNA5SP342 | N-above, 0.07 | Ribosomal pseudogene 342 |
| HSPA1L | N-above, 0.05 | Expressed in normal prostate epithelium as part of the cellular stress response machinery [47]. It helps maintain protein homeostasis and cell survival under stress. More experimental evidence is needed. |
| **Perfect panel of T-genes** | **Type and Frequency in tumor samples** | **Reported oncogenic role** |
| EPHA10 | T-above, 0.74 | Oncogenic driver; m6A-methylation regulated by |
|  |  | RBM15B; activates ERK/AKT pathway [48]; potential immune evasion [49,50]. |
| UCN | T-above, 0.64 | UCN-CRFR2 signaling may act as an endogenous inhibitor of angiogenesis and proliferation; receptor loss (CRFR2) may disrupt tumor suppression [51,52]. |
| TMEM86A | T-above, 0.51 | Transmembrane protein; epigenetically regulated (H3K27me3) [53]; cited in prostate cancer context. |
| RPL26P23 | T-above, 0.43 | No information found; Pseudogene (likely) |
| RBM15B | T-above, 0.41 | Oncogenic m6A writer; Methylates EPHA10, PCNA mRNAs; stabilizes transcripts; promotes proliferation [54]. |
| NPY4R2 | T-above, 0.24 | A member of the neuropeptide Y receptor family. Other neuropeptide receptors (e.g., NPY1R, NPY5R) have been implicated in prostate cancer progression, neuroendocrine differentiation, and bone metastasis. By analogy, NPY4R2 could play a role, but no data supports this speculation [55]. |
| TEKT4 | T-above, 0.24 | Chemotherapy resistance; Microtubule-associated protein; mutations alter taxane binding [56]; linked to ERG fusion status [57]. |
| ENSG00000271895 | T-above, 0.20 | No information found |

### 3d. Labeling the somatic evolution of a normal tissue with the help of perfect gene panels

In this section, we present a “taxonomy of the normal tissue,” i.e., we label its somatic evolution state. We use the idea of perfect panels of N-genes, described in Ref. 6. Any normal tissue sample is characterized by at least one activated N-gene in the panel. Note that, in principle, the perfect panel is not unique; we may find different sets that achieve perfect classification. Again, we use TCGA data for PRAD as an example.

We start from the gene with the highest activation frequency in the set of normal samples, *SEPTIN10*, and add genes until all samples in the set show activation of at least one gene. The procedure is detailed in Ref. 6. One realization of this procedure leads to the following panel of 11 genes: {*SEPTIN10, CA14, TES, PARM1, APOBEC3C, ENSG00000224984, TMT1A, SLC18A2, SNORD104, RNA5SP342, HSPA1L*}. Their frequencies and properties are shown in Table I. Note that all of them, except *SNORD104*, belong to the N-above category, i.e., they are tumor suppressor–like genes. In addition, five of them are NT genes: *SEPTIN10, TES, PARM1, APOBEC3C*, and *SNORD104*. This means they can switch their role from functioning as a brake in the normal state to boosting tumor progression.

The somatic evolution stage of a normal tissue may be labeled, as seen in previous sections, by the total number of activated N-markers. Additionally, we can indicate which pathways or mechanisms are protecting the normal tissue from becoming a tumor. Panel genes are indicators of the active mechanisms. Unfortunately, the gene ontology and biological function of many genes are, at present, unknown or only partially known. We list in Table I some of the known properties of these genes, particularly those related to the protective role they play. The table shows that the panel genes constitute a meaningful set. There is no information about the novel transcript *ENSG00000224984* or the ribosomal pseudogene *RNA5SP342*.

In Figure 4a we show the classification that panel genes induce in the set of normal samples. A cell with a black color means that the gene marker is active. Samples are listed according to the number of total active N-genes. That is, less evolved samples with a larger number of N-genes appear at the bottom, and samples closer to becoming a tumor appear at the top of the figure.

**Figure 4.**
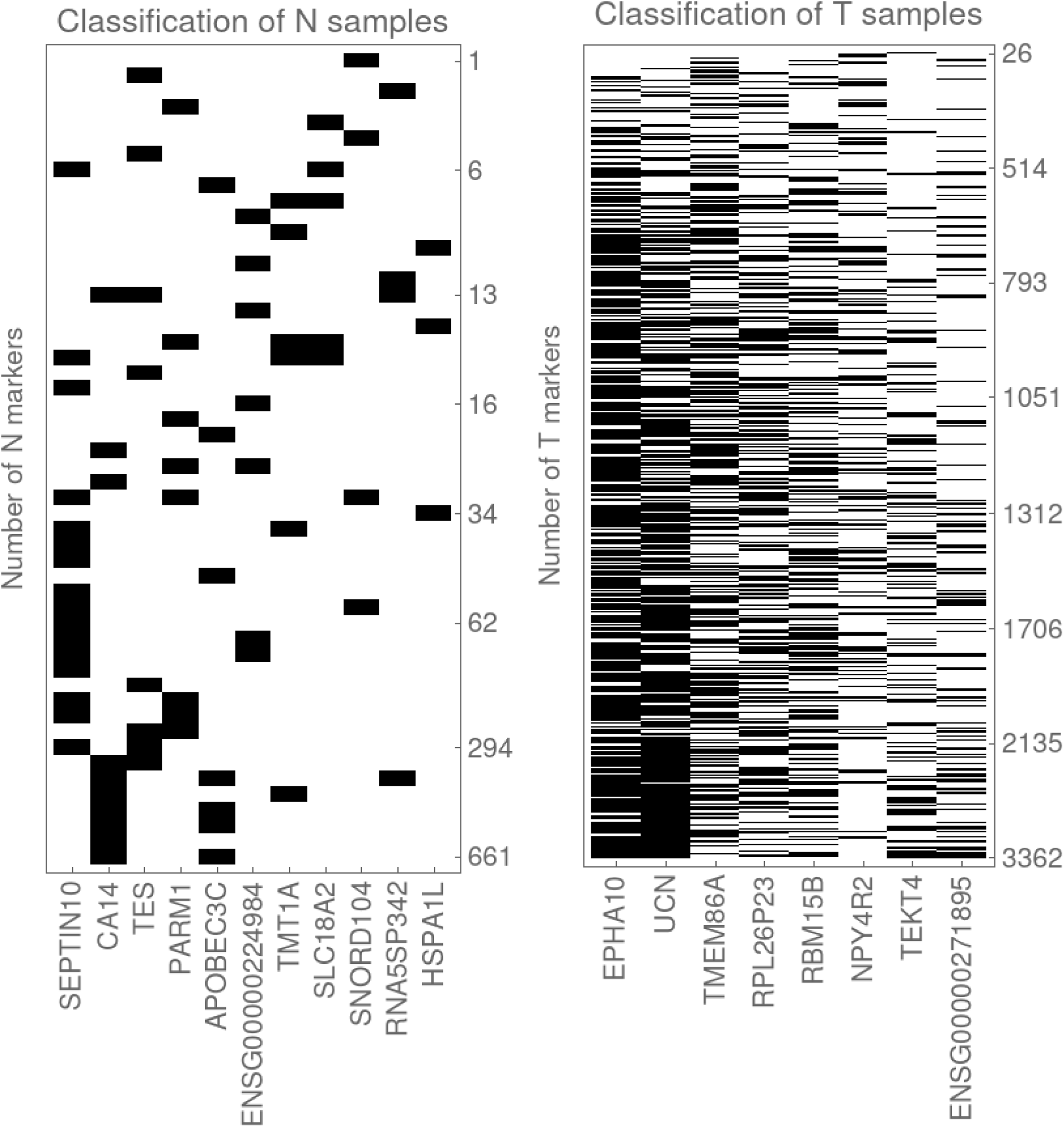
Taxonomy of normal tissues and tumors suggested by the perfect panels. a) The 52 normal tissue samples in PRAD (rows in the array plot) are classified by the number of active N-markers (evolution stage) and by which genes in the perfect panel of 11 N-genes are active (black cells in the plot indicate active protection mechanisms). b) Classification of the 499 tumor samples by the number of active T-markers and active panel genes. The latter are related to the mechanisms that boost tumor progression.

Notice that the classification is perfect. Any normal sample shows between one and three activated panel genes. It seems that there is a weak correlation between the number of activated protection mechanisms and the degree of somatic evolution. Healthy samples show two or one, rarely three, activated genes, whereas evolved samples show, as a rule, only one. The small number of activated genes probably indicates that the protection mechanisms are inherited. More data are needed to confirm this hypothesis.

We may use the patterns of active panel genes to cluster the samples. Figure 5 shows the dendrogram resulting from applying a Dice dissimilarity distance for this purpose. With a threshold distance of 0.7, five groups and a single isolated pattern ( *HSPA1L* activated) are obtained. Of course, this is the first time a molecular classification of normal samples has been proposed, and its clinical relevance needs to be validated.

**Figure 5.**
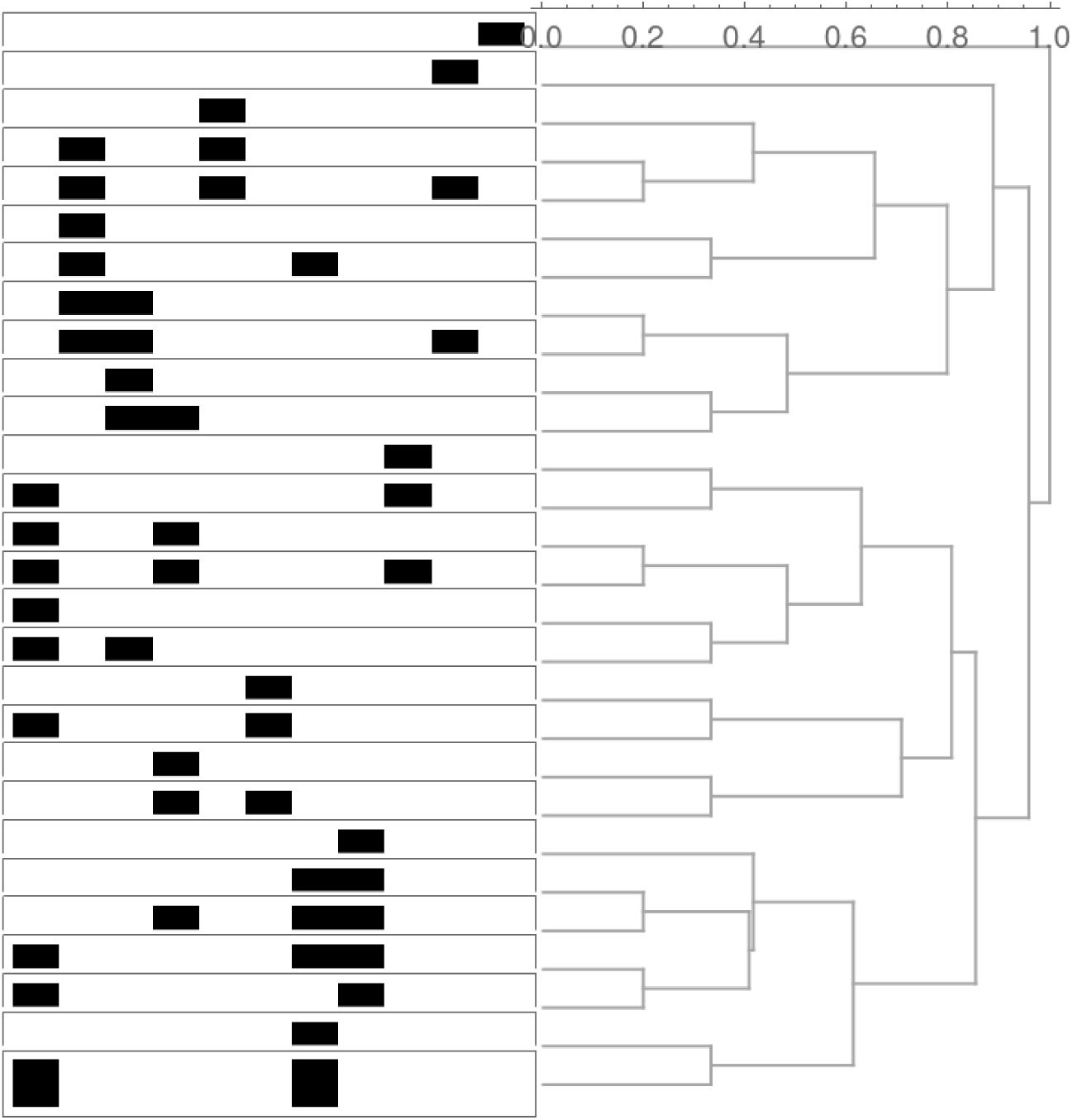
Clustering of normal samples according to a Dice dissimilarity distance. There are 28 distinct patterns for the combinations of panel genes. A threshold distance of 0.7, for example, defines five clusters and a single independent pattern (only HSPA1L active).

Let us stress that panel genes are not block genes. The only exception is *HSPA1L*, which is a representative of four genes with the same expression profile: {*HSPA1L, HSPA1A, BAG3, DNAJB1*}, all related to the stress survival network and the stabilization of the androgen receptor.

### 3e. Labeling the clonal evolution of tumors with the help of perfect gene panels

Similarly, we may use a perfect panel of T-genes to achieve perfect classification of tumors. An activated gene in the panel would indicate a pathway or mechanism that facilitates evolution toward a more malignant state. Proceeding along the lines of Ref. 6, we obtain the following perfect panel of 8 genes: {*EPHA10, UCN, TMEM86A, RPL26P23,*

*RBM15B, NPY4R2, TEKT4, and ENSG00000271895*}. All of these genes belong to the T-above group; thus, they are oncogene-like. Their frequencies in the set of tumor samples and their known oncogenic attributes are listed at the end of Table I, also evidencing that the set is meaningful. The genes *RPL26P23* and *ENSG00000271895* are not annotated; their properties are completely unknown.

In Figure 4b we show how the use of the number of active T-genes and active boosting mechanisms, labeled by the panel genes, provides a complete classification of tumor samples. Recently formed tumors exhibit one or two active panel genes, whereas evolved tumors show, as a rule, six. In the same way we did for normal samples, the patterns of active mechanisms may be used to group or cluster the tumor samples. The corresponding dendrogram is shown in Figure 6.

**Figure 6.**
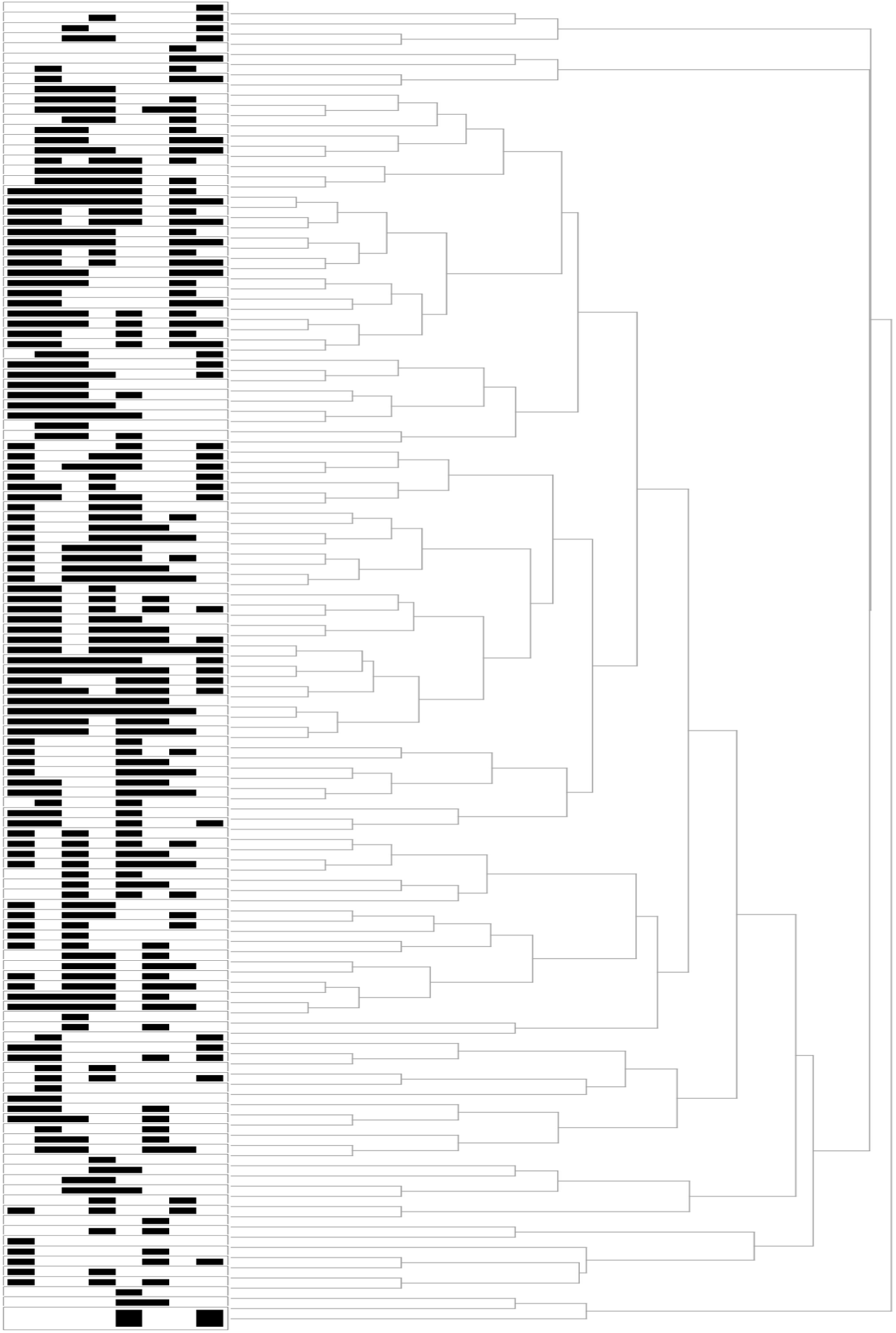
Clustering of tumor samples. There are 129 independent combinations of panel genes, which can be clustered by selecting an appropriate threshold distance.

A detailed discussion of the clinical relevance of the proposed classification is beyond the scope of the present paper. In the next section, we nevertheless briefly discuss the connection with the standard taxonomy of prostate tumors, based on mutational drivers.

### 3f. Comparison with traditional driver genes in PRAD

There exist different molecular classifications of prostate tumors [58]. As an example, we briefly revisit the TCGA classification, which is essentially multi-omics but is labeled by a set of seven mutational driver genes [59], listed in Supplementary Table I.

The classification algorithm was applied to an initial cohort of 333 samples, not the 499 tumor samples we are using in our study. The obtained panel of genes is incomplete, offering a classification for 76% of the samples. In Supplementary Table II, we show a schematic representation of the TCGA classification, which illustrates the principle guiding the search for panel genes: their mutually exclusive character. In other words, when a gene is mutated in a sample, the other genes in the panel are not mutated.

In contrast, our classification is complete, including 100% of the samples, and the number of active panel genes is an indicator of the degree of clonal evolution of the tumor.

Let us examine the TCGA panel genes in light of our scheme. Four of the genes— *ERG, ETV1, ETV4*, and *FLI1*—belong to the ETS family of transcription factors and show characteristic fusion patterns in PRAD. The other three genes—*SPOP, FOXA1*, and *IDH1* —exhibit characteristic mutations in PRAD. Note that they consider fusion or mutation frequencies as low as 1%.

Recall that in our analysis, a threshold of 10% for the deregulation frequency of T-genes, dictated by the Fisher test, is included. If we relax this condition, the four fusion genes in the panel and *IDH1* may be classified as T-outside genes, whereas *SPOP* and *FOXA1* are included in the NT group. It is with this relaxed view that the seven TCGA panel genes become meaningful in our description.

With regard to mutations, they are irreversible changes. Due to the aforementioned higher entropy of tumors, mutations are in general more compatible with tumors than with the normal state. The regulation networks deal with expression values. Mutations are like nodes in the network with forced activation. A mutation in an NT gene in the normal state reversing its activation to tumor-like constitutes a forced T-marker that may accelerate tumorigenesis. The two NT genes in principle have this potential, although their observed frequencies in the tumor subset are low. On the other hand, a relevant mutation in any of the other five genes is a forced marker boosting tumor progression.

In general, the frequency of observed relevant mutations in PRAD is low [60]. In contrast, expression deregulations are very common, reaching a maximum frequency of around 74% for the *EPHA10* gene in the set of tumor samples. The histogram showing the number of T-genes in PRAD as a function of their deregulation frequencies is presented in Supplementary Fig. 3. There are more than 200 genes deregulated in 30% of tumor samples, for example. In other tumors, such as LUAD and LIHC, deregulation frequencies are even higher. This is another advantage of our scheme.

## 4. Concluding remarks

We have stressed the significance of N- and T-genes in the process of carcinogenesis and the clonal evolution of tumors.

The normal tissue starts with a set of N-markers, which are partially deactivated in the course of somatic evolution as a result of subsequent deregulation cascades. Thus, the activated N-genes in a given normal sample are in fact markers and anchors, and when deactivated, they delineate the pathway or trend toward the tumor. The number of active N-markers defines the stage of somatic evolution of the tissue, whereas the mechanisms still protecting the tissue may be labeled by the active markers in a perfect panel of 11 genes.

On the other hand, cascades in a formed tumor continuously activate T-markers. The number of activated markers defines the stage of clonal evolution of the tumor. The mechanisms facilitating evolution toward a more malignant state may be labeled by the active markers in a perfect panel of 8 genes.

The classification schemes for normal tissues and tumors derived from our perfect panels need to be clinically validated.

Many interesting questions remain to be answered. The most important of these is perhaps the proper description of the dynamics. To this end, we need the regulation networks. In Ref. [61], we construct probabilistic deregulation networks for N- and T-genes. These are inferred directly from the gene expression data. As mentioned above, both networks should be included in the description of the transition point, where the NT-genes could play a decisive role. Research along some of these directions is currently in progress.

## Supplementary Material

**Supplementary Figure 1.**
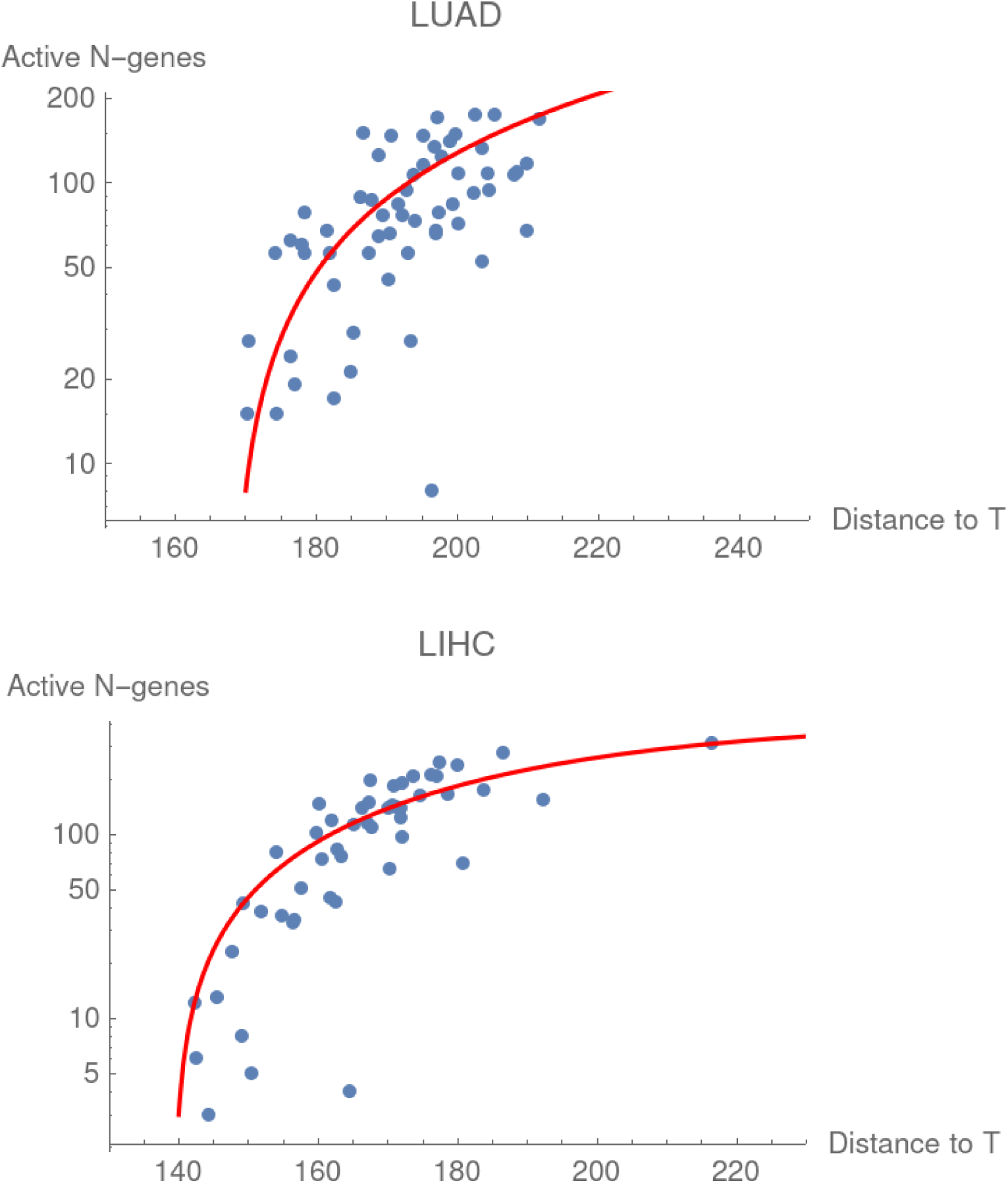
Results similar to Fig. 3a, that is number of N-genes as a function of the distance to the tumor attractor, for lung adenocarcinoma (LUAD) and liver hepatocarcinoma (LIHC).

**Supplementary Figure 2.**
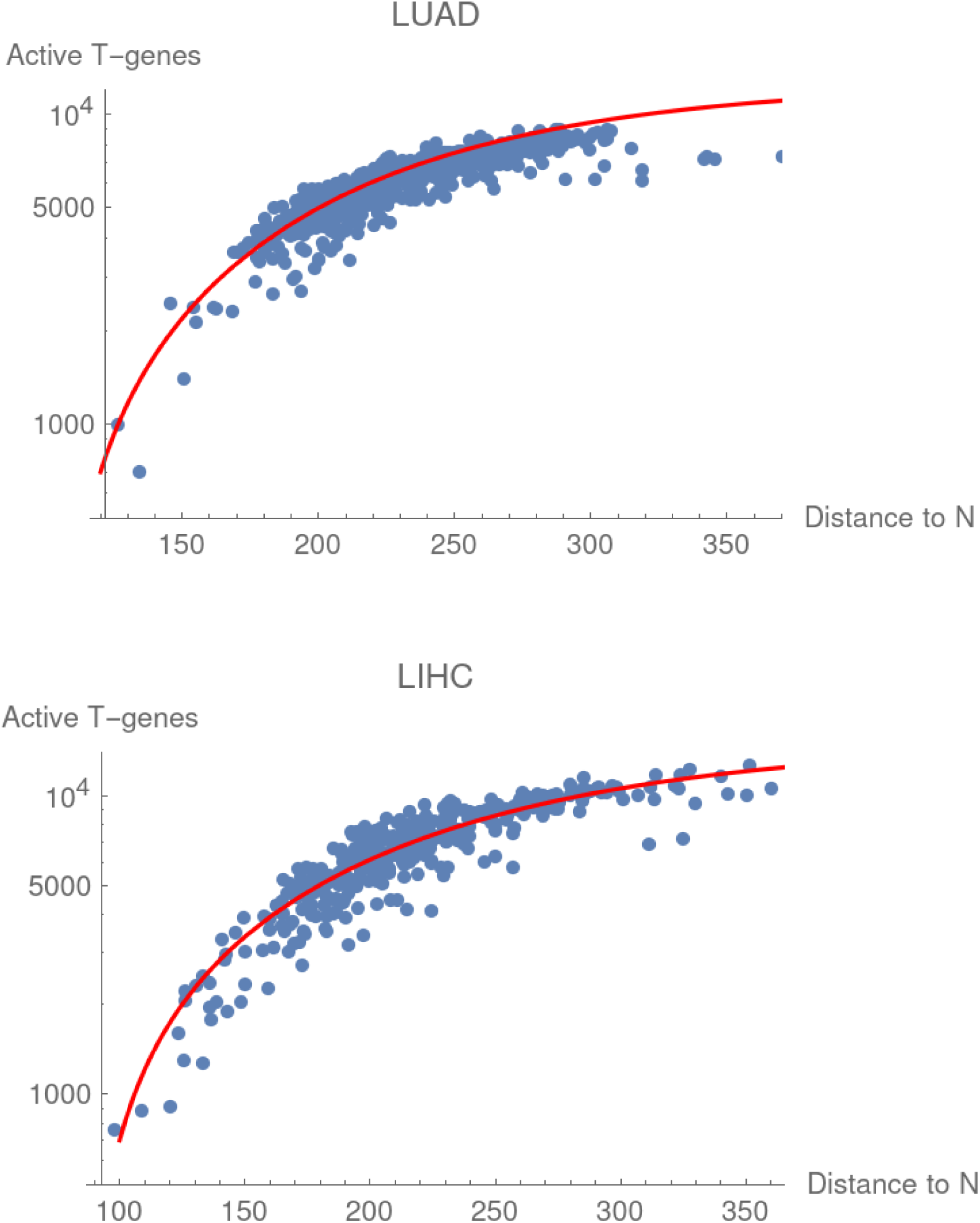
Number of active T-genes as a function of the distance to the N-attractor in LUAD and LIHC, respectively.

**Supplementary Table I.** Mutational drivers used in the TCGA classification of PRAD tumors. The Freq(+) value is the fraction of samples in the top interval. For a T-outside gene, this is a tumor-only interval; however, for the SPOP gene, it is a normal-only interval that includes 15.4% of normal samples. Accordingly, the Freq(-) value is the fraction of samples in the bottom interval.

| Gene | Mutation or Fusion Freq. | Classif. | Freq(+) | Freq(-) |
| --- | --- | --- | --- | --- |
| ERG (fusion) | 0.46 | T-outside? | 0.23 | 0.01 |
| ETV1 (fusion) | 0.08 | T-outside? | 0.08 | 0.08 |
| ETV4 (fusion) | 0.04 | T-outside? | 0.05 | 0.05 |
| FLI1 (fusion) | 0.01 | T-outside? | 0.01 | 0.06 |
| SPOP (mutation) | 0.11 | NT-gene? | 0.154 (N) | 0.042 (T) |
| FOXA1 (mutation) | 0.03 | NT-gene? | 0.046 (T) | 0.154 (N) |
| IDH1 (mutation) | 0.01 | T-outside? | 0.04 | 0.02 |

**Supplementary Table II.** Schematics of the TCGA classification of PRAD tumors.

| Samples → | (46%) | (8%) | (4%) | (1%) | (11%) | (3%) | (1%) | (26%) |
| --- | --- | --- | --- | --- | --- | --- | --- | --- |
| ERG | 1 | 0 | 0 | 0 | 0 | 0 | 0 | 0 |
| ETV1 | 0 | 1 | 0 | 0 | 0 | 0 | 0 | 0 |
| ETV4 | 0 | 0 | 1 | 0 | 0 | 0 | 0 | 0 |
| FLI1 | 0 | 0 | 0 | 1 | 0 | 0 | 0 | 0 |
| SPOP | 0 | 0 | 0 | 0 | 1 | 0 | 0 | 0 |
| FOXA1 | 0 | 0 | 0 | 0 | 0 | 1 | 0 | 0 |
| IDH1 | 0 | 0 | 0 | 0 | 0 | 0 | 1 | 0 |

**Supplementary Fig. 3.**
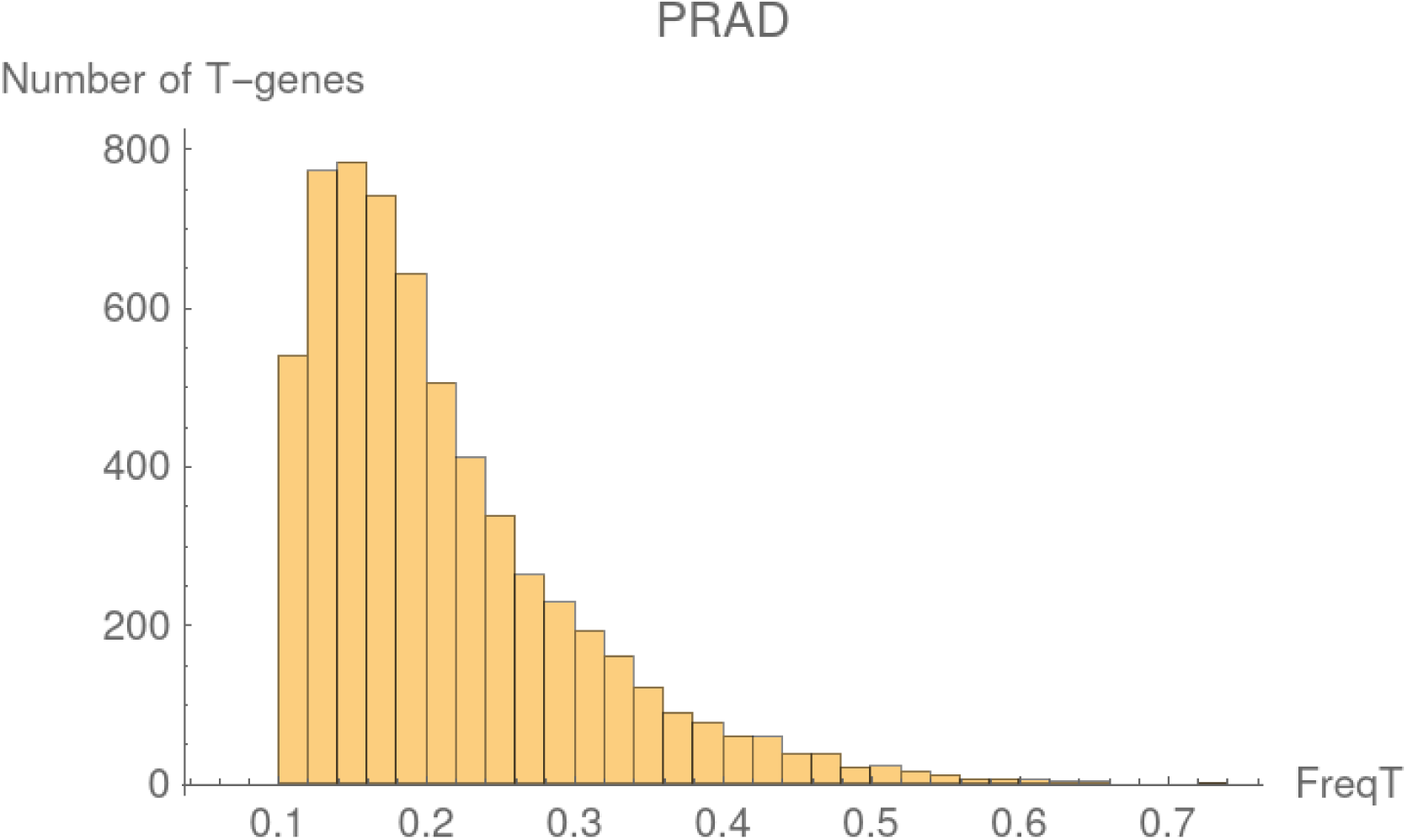
The number of T-genes as a function of their deregulation frequency in PRAD.

